# A CRISPR-Cas9 assisted analysis of single-cell microbiomes for identifying rare bacterial taxa in phycospheres of diatoms

**DOI:** 10.1101/2024.11.10.622248

**Authors:** Ruben Schulte-Hillen, Jakob K. Giesler, Thomas Mock, Nigel Belshaw, Uwe John, Tilmann Harder, Nancy Kühne, Stefan Neuhaus, Sylke Wohlrab

## Abstract

Primary production in aquatic systems is governed by interactions between microalgae and their associated bacteria. Most of our knowledge about algal microbiomes stems from natural mixed communities or isolated algal monocultures, which therefore does neither address the role of genotypic diversity among the algal host cells nor do they reveal how this host diversity impacts the assembly process of associated bacteria. To overcome this knowledge gap, we developed a single-cell 16S sequencing approach in combination with CRISPR-Cas9 guided depletion of host 16S contaminations from the chloroplast. The validity of this novel method was tested by comparing bacterial communities of 144 single-cells across three genotypes of the Arctic marine diatom *Thalassiosira gravida* grown under different environmental conditions. From these, 62 single-cells were additionally sequenced after CRISPR-Cas9 treatment. Due to the improved sequencing depth, bacterial richness associated with individual diatom cells was increased by up to 56%. By applying this CRISPR-Cas9 treatment we not only revealed intraspecific host-genotype associations but also rare bacterial taxa that were not detected by standard 16S rRNA gene metabarcoding. Thus, the CRISPR-Cas9 assisted single-cell approach developed in this study advances our understanding on how the intraspecific diversity among algal hosts impacts the assembly process of their associated bacteria. This knowledge is essential to understand the co-evolution and adaptation of species in algal microbiomes.

## Introduction

The role and function of interactions between individuals in microbial communities is increasingly recognized in microbiome research (Margulis, 1991, Dittami et al., 2021), which coined the term ‘holobiont’ describing an assemblage of a host (e.g., microalga) and associated microbes (e.g., bacteria). As the main primary producers in aquatic ecosystems, both uni- and multicellular algae have been studied from a holobiont perspective because their associated microbiomes substantially affect algal physiology, growth and resilience (Amin et al., 2012, Giesler et al., 2023). For example, interactions between host algae and bacteria are underpinned by specific bacterial enzymes as well as reciprocal nutritional requirements for essential trace elements, micro-, and macronutrients (Amin et al., 2012, Kodama and Fujishima, 2022).

Given the recognized importance of microalgae for diverse ecosystem functions (Cirri and Pohnert, 2019, Kuhlisch et al., 2023) methods to accurately determine their associated microbial communities are of considerable value. Current strategies to assess the bacterial composition associated with microalgae predominantly rely on laboratory monocultures or natural field communities, introducing uncertainties such as culture artifacts and biases due to single-cell isolation and filtration. Moreover, clonality in microalgal populations usually is short-lived as sub-populations arise due to genetic drift leading to clonal interference. How this intraspecific host diversity impacts the diversity of associated bacterial communities has rarely been studied and is therefore largely unknown. Natural algal microbiomes, on the other hand, are difficult to study in a natural community context due to challenges in assigning bacteria to specific hosts for revealing their interactions (Martin et al., 2021). To address these challenges, single-cell-based metabarcoding has recently been developed to assesses the microbial diversity associated with single-cells of host protists such as choanoflagellates and ciliates (Needham et al., 2022, Boscaro et al., 2023, Rossi et al., 2019). However, this single-cell sequencing strategy comes with several challenges. For instance, sequencing small amounts of DNA isolated from a proximity of the host protist might introduce a bias due to significant variability between single host cells in terms of the success of DNA extraction and amplification (Schmitz et al., 2021). Although microbiomes on single sand grains have been successfully studied despite low DNA concentrations (Probandt et al., 2018), similar microbiome analyses using single-celled algal microbiomes have been challenging due to the presence of 16S rRNA gene copies encoded in the algal chloroplast genomes, which can account for up to 99% of all sequence reads (Lundberg et al., 2012, Zarraonaindia et al., 2015).

This predominance of 16S rRNA gene copies encoded in chloroplast genomes reduces the sequencing depth of the 16S copies from the associated microbiomes. This caveat is adding to challenges imposed by overall low quantities of DNA, spatial heterogeneity and potential other methodological biases (see above). One strategy to reduce contamination by host 16S rRNA gene sequences from chloroplasts is PCR clamping, a method to suppress a particular allele during PCR (Orum et al., 1993). Yet, this strategy has its own biases, i.e., species-specific suppression of PCR amplification (Jackrel et al., 2017, Baker and Kemp, 2014). Here, we address these biases with a novel approach, **S**ingle-**C**ell **Ho**lobiont **C**as9-**O**ptimised-**Seq**uencing (SCHoCO-Seq), which employs a Cas9 nuclease together with a target-specific guide RNA to dissect the 16S rRNA gene sequences of oceanic microalgae. Our approach is thus methodologically complementary to the approach developed by Song and Xie (2020), who used Cas9 to dissect the 16sRNA gene copies of rice chloroplasts from bulk leaf samples. However, we further extended this method to the single-cell level. In addition, as our approach is not restricted to a single species, it can be used to gain insight into a wide range of natural microbiomes associated with diverse single-cell host organisms.

We have compared this novel SCHoCO-seq approach with CRISPR/Cas9 untreated single-cell control microbiomes (further referred to as SC-Seq) of the diatom *Thalassiosira gravida* under different culture conditions. This workflow of SCHoCO-seq is based on single-cell isolation, optional DNA extraction, subsequent nested low template PCR and Cas9-mediated, target-specific cleavage of the diatom chloroplast 16S rRNA gene amplicons, followed by amplicon metabarcoding of the associated prokaryotic community. Furthermore, controls were conducted to test for contaminations and off-target activity.

## Material and methods

Three strains of the Arctic diatom *T. gravida* were grown under different culture conditions. Single-cells were manually isolated and washed to remove free-living bacteria leaving mainly diatom-associated bacteria. The isolated single diatom cells were processed according to the workflow in Figure 1 and described in detail below. Two different pre-processing protocols are presented: a direct PCR method, which is less labor intensive, and a method involving a single-cell DNA extraction step, which has the advantage of retaining the DNA for additional analysis.

**Figure 1:**
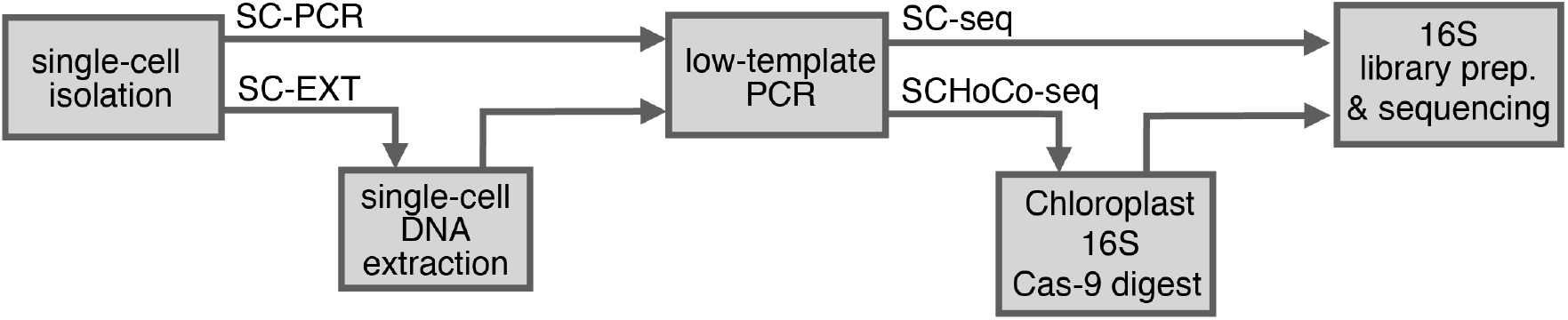
Schematic overview of the laboratory workflow. After single-cell isolation, samples were either subjected to DNA extraction (SC-EXT) or processed directly to low template PCR without prior DNA extraction (SC-PCR). A subset of low-template PCR products from all samples (62 out of 144) were treated with Cas9 to dissect the chloroplast 16S rRNA gene (SCHoCO-seq) prior to library preparation.

### Culture Conditions

Diatom cultures of *Thalassiosira gravida* strains (UIO 478; UIO 483; UIO 448, table S1) were obtained from the Norwegian Culture Collection of Algae (NORCCA). Here, we refer to UIO 478 as A1, UIO 483 as A2, and UIO 448 as A5. Stock cultures were maintained in three culture media based on a modified and silica-enriched K-medium (20) (table S2), a nitrogen-deplete medium (table S3), and a vitamin-deplete medium (table S4). Cultures were grown at 4°C and 50 μmol photons m^-2^ s^-1^ under a light:dark period of 16:8 h.

### Single-cell Isolation

To study the diversity of diatom-associated microbiomes among conspecific host cells across genotypes, single-cells of *T. gravida* strains A1, A2 and A5 cultured under different conditions (see above) were isolated (n = 8). To remove the free-living bacteria, Individual diatom cells were picked and sequentially washed five times by pipetting 3 μL into two separate 30 μL drops of PBS buffer (PBS CellPure, Roth, Karlsruhe, Germany), followed by three washings in 30 μL of PCR-grade water under an inverted light microscope. To validate the absence of free-living bacteria, 3 μL culture blanks (without diatom cells) were sampled from all strains and treated identical to the single-cell samples undergoing single-cell DNA extraction (see below). Cells were stored either in 30 μL 2xT&C lysis buffer of the MasterPure Complete DNA & RNA Purification Kit (Lucigen, Wisconsin, USA) or in 10 μL PCR-grade water. Cells in PCR-grade water were used directly as DNA template for a nested, low template PCR (see below). Cells stored in 2xT&C lysis buffer were used to generate a PCR product after DNA extraction (see below).

### DNA Extraction from Isolated Single-cells (SC-EXT)

To extract DNA from the single-cellular diatom microbiomes, a modified protocol of the DNA MasterPure Complete DNA & RNA Purification Kit was employed (SC-EXT). The changes to the manufacturer instructions were: (i) smaller buffer volumes, (ii) added pellet dye, and (iii) rinsing of pipette tips. (details in table S5)

### Nested Low Template PCR of Single Diatom Cell Holobionts

During the low template PCR, the hypervariable V1-V9 region of the 16S rRNA gene was targeted by Primer 27F (5’-AGAGTTTGATYMTGGCTCAG-3’) (Frank et al., 2008) and 1492R (5’-TACGGYTACCTTGTTACGACTT-3’) (Lane D, 1991). PCR reactions for single-cell samples were set up to 50 μL per reaction containing 5 μL 10 x Hotmaster Taq Buffer (Qiagen, Hilden, Germany), 0.5 μL Primer 27F (10 mM), 0.5 μL Primer 1492R (10 mM), 0.5 μL dNTPs (10 mM), 0.1 μL bovine serum albumin (BSA, 10 mg/mL), 0.5 Taq Polymerase (Qiagen), 32.9 μL H_2_O and 10 μL Single-Cell Sample (SC-PCR) or 2.5 μL extracted DNA (SC-EXT) plus 7.5 μL H_2_O. PCR protocols for SC-PCR and SC-EXT differed only in the duration of the initial denaturation phase of 94°C for 1.5 min (SC-EXT) and 10 min (SC-PCR), respectively. Both protocols consisted of 35 cycles; denaturation: 94°C for 1.5 min primer annealing: 55°C for 1.5 min, elongation: 68°C for 2 min, final elongation: 68°C for 10 min. PCR products were cleaned with the MinElute PCR Purification kit (Qiagen). All culture blanks (without diatom cell) showed no bands during the gel-electrophoresis (Fig. S1).

### Cas9 digest of chloroplast-derived 16S-rRNA gene amplicons

The 16S rRNA gene amplicons generated by nested low template PCR were cloned into pCR 4-TOPO vectors and transferred into competent *E. coli* cells of the TOPO TA cloning kit (Thermo Fisher, Massachusetts, USA). Plasmids of single colonies were purified with the Plasmid Plus Midi Sample Kit (Qiagen). PCR products were sequenced on an ABI 3130xl Genetic Analyzer; obtained sequences were searched for chloroplast 16S rRNA gene sequences by BLAST search in the PhytoRef database (Decelle et al., 2015). Plasmids containing the respective *Thalassiosira* chloroplast 16S rRNA gene and respective sequences were used for the design of the gRNA.

### Design of gRNA

Guide RNAs (gRNAs) were designed for the *T. gravida* chloroplast 16S rRNA gene according to Hopes et al. (2017). Five gRNA sequences were selected for in vitro testing by comparison with endosymbiotic derived 16S rRNA gene amplicons and gRNAs lacking homology in the PAM sequences and therefore not cut by the gRNA mediated Cas9 (Hsu et al., 2013). Subsequently, the selected gRNAs were tested for their efficiency in directing Cas9 digestion of the *T. gravida* chloroplast 16S rRNA gene by OmicronCR, Norwich, UK.

### Selection of the most suitable gRNA

The five gRNA sequences were aligned to the chloroplast sequence of a total of 15 diatom families retrieved from the PhytoREF plastidal 16S rRNA database (Decelle et al., 2015). The gRNA which cut the central V4 region of chloroplast ASV-16S RNA sequences in a highly conserved region was selected (Fig. S2). To determine the general utility of SCHoCO-seq for diverse diatom taxa, we searched for matching crRNA target sequences in the full PhytoREF database (Decelle et al., 2015). In addition, we searched the SILVA database V138 (Quast et al., 2013) for potential off-target sequences to further test whether a loss of microbial 16S rRNAs occurred due to unwanted Cas9 digestions. We allowed two mismatches to include potential off-targets with a 4% probability given the low mismatch tolerance of CRISPR-Cas9 (Anderson et al., 2015). Furthermore, we evaluated in-silico the phylogenetic range of the diatom chloroplast sequences not only regarding matching gRNA, but also regarding the outer and inner primers of the SCHoCO-seq approach. For this purpose, we also relied on the PhytoREF database. Chloroplast sequences with 100% matches to all primers and the gRNA were aligned with kalign (Lassmann, 2019) and an unrooted phylogenetic tree was built with FastTree (Price et al., 2009). Taxonomic annotations were transferred from the PhytoREF database (Decelle et al., 2015).

### Synthesis of sgRNA

The single guide RNA (sgRNA) comprised two specific sequences: the CRISPR RNA (crRNA) containing the complementary sequence to the target sequence, and the trans-activating-crRNA (tracrRNA) guiding and stabilizing the Cas9 nuclease (Hiranniramol et al., 2020). Both parts were derived from respective oligonucleotides (universal oligo: 5’-AAAAAAGCACCGACTCGGTGCCACTTTTTCAAGTTGATAACGGACTAGCCTTATTTTAACTTGCTATTTCT-3’, specific oligo: 5’-TAATACGACTCACTATAGGAAGTCAACTGTTAAATCTTGGTTTTAGAGCTAGAAATAG-3’) (Eurofins, Hamburg, Germany) which were grafted together and amplified in a PCR forming the double-stranded sgRNA template. For this reaction to succeed the two oligonucleotides must share a common overlapping sequence (linker) as well as binding sequences for a forward and reverse primer (Fig S2).

PCR was set up to 50 μL, containing 5 x Phusion Plus buffer (10 μL), a sgRNA specific oligo (1 μL, 0.1 μM), T7 sgRNA oligo 2 (1 μL, 0.1 μM), T7sgRNA Forward (3.75 μL, 10 μM), T7sgRNA Reverse (3.75 μL, 10 μM), dNTPs (1.5 μL 10 μM), Phusion polymerase (0.5 μL) and PCR grade water (28.5 μL) (all Jena Bioscience, Germany). PCR conditions were: initial denaturing (98°C, 30 s), 30 cycles of denaturing (98°C, 10 s), annealing (51°C, 10 s), and elongation (72°C, 15 s); final elongation (72°C, 10 min.). The sgRNA was synthesized by *in-vitro* transcription as followed: 20 μL containing High Yield T7 reaction buffer (2 μL), DTT (2 μL, 100 mM), ATP, GTP, CTP, UTP (each 1.5 μL, 100 mM), High Yield T7 RNA Polymerase Mix (2 μL), template DNA (2 μL) and PCR grade water (4 μL) (Jena Bioscience, Germany). The reaction mix was incubated for 11 h at 37.5 °C, followed by a DNAse I digest (1 μL RNAse free DNAse I, Jena Bioscience) for 15 min at 37°C. The sgRNA was purified using the RNA clean & concentrator-5 kit (Zymo Research, USA) and the gRNA synthesis was checked for correct product size with an RNA Nano Chip Assay on a 2100 Bioanalyzer device (Agilent Technologies, Santa Clara, California, USA).

### In-vitro digestion of 16S amplicons using Cas9 and gRNA

The Cas9 digest was performed after the amplification and purification of the V1-V9 region of the prokaryotic 16S rRNA and prior to the 16s rRNA V4 amplicon sequencing library preparation. The Cas9 Nuclease (New England Biolabs, USA) and the gRNA were assembled in a 27 μL reaction containing PCR grade Water (20 μL), Cas9 Buffer (3.5 μL), gRNA (150 nM, 3 μL) and Cas9 (1 μM, 0.5 μL) by incubating at 25°C for 10 min. The 16S rRNA gene was digested by adding 3 μL of 15 nM target DNA to the mixture and incubating it at 37°C overnight (∼11 h).

The sample was cleaned with the MinElute PCR Purification kit (Qiagen).

### Illumina Sequencing of partial 16S rRNA genes

The amplicon library was constructed according to the manufacturer’s protocol for 16S metagenomic sequencing library preparation, described in Ahme et al. (2023). The library was sequenced on the MiSeq System (Illumina, San Diego, USA) according to the MiSeq Reagent Kit v3 (600 cycles) (Illumina). Forward Primer: MS_v4_515F_N: 5’-TCGTCGGCAGCGTCAGATGTGTATAAGAGACAG+GTGCCAGCMGCCGCGGTAA-3’ (Apprill et al., 2015), Reverse Primer: MS_v4_806R_1: 5’-GTCGTCGGCAGCGTCAGATGTGTATAAGAGACAG+GGACTACHVGGGTWTCTAAT-3’ (Parada et al., 2016).

### Processing of sequence data

FASTQ files were demultiplexed according to the ‘Generate FASTQ workflow’ of the MiSeq sequencer software. Primers were removed with cutadapt v2.8 (Martin, 2011), and sequence data were processed with the DADA2 R package (quality trimming, denoising, merging, removal of chimeras, Callahan et al., 2016). The resulting Amplicon Sequence Variants (ASVs) were taxonomically annotated with the SILVA v138 database (Quast et al., 2013). Details about the FASTQ and ASV data processing pipeline are described in Ahme et al. (2023). Statistical analyses were performed in R version 4.1.2 (R Studio, Inc. 2021; www.r-project.org).

### Analysis of single-cell diversity and bacterial community composition

ASVs were excluded from a sample if 1) their relative contribution to the respective sample was smaller than 0.25% as recommended by Reitmeier et al (2021) 2) ASVs represent common contaminants according to Sheik et al. (2018) 3) ASVs were known to be derived from non-marine sources (see supplemental table S6). Samples with sequencing depths contained in the lower 10% quantile of the whole dataset, or below 10000 reads, were excluded from the analysis. ASV count tables were scaled by subsampling using the SRS package (Beule and Karlovsky, 2020) and normalized by power transformation (X^0.25^). Graphical representation was done with the Phyloseq package (McMurdie and Holmes, 2013).

### Statistical analysis of single-cell samples

Richness and Shannon diversity of the *T. gravida* single-cell samples were determined using the “microbiome” package (Lahti, 2017). To test for significant effects on ASV richness and Shannon diversity as a response to strain identity (A1; A2; A5), culture condition (full medium; -N; -Vit) and DNA processing method (SC-EXT; SC-PCR), three-way ANOVAs were calculated with subsequent Tukey’s post-hoc tests. To evaluate differences in microbiome community composition across *T. gravida* strains and culture conditions, a Bray-Curtis distance matrix was calculated from the down-sampled and normalized ASV count data, serving as input for ordination of a principal component analysis. PERMANAOVA was applied on the Bray-Curtis distance matrix as dependent variable, and strain identity and culture condition as independent variables using the “adonis2” function of the “vegan” package (Oksanen et al., 2013). To obtain a more detailed analysis on microbiome composition and its drivers, bacterial ASVs were aggregated and subset to the genus level. Normalized count subsets for each genus were tested for differences across strains and culture conditions using two-way ANOVAs.

### Analysis of SCHoCO-seq samples

Bacterial ASV data of single-cells of *T. gravida* strains A1, A2 and A5 were quality-filtered, down-sampled and normalized as described above. After richness calculation, an ANOVA was conducted to test for differences in bacterial ASV richness as a response to SCHoCO-seq treatment (compared to SC-seq control treatment). The respective diatom single-cell ID (of which the DNA product for both treatments originated) was treated as a random factor. To test for differences in relative proportions of bacterial ASVs, chloroplast ASVs, and mitochondrial ASVs as a response to SCHoCO-seq, a t-test was conducted to compare the relative proportion of each ASV category compared to the control treatment. To evaluate the effect of sequencing depth on the relative proportion of bacterial, chloroplast, and mitochondrial ASVs, 1000 replicates of random subsamples (each containing 500, 1000, 5000, and 10000 ASVs) were generated from each sample.

The incidence rate ratio (IRR) represents the probability of detecting a bacterial family with SCHoCO-seq compared to SC-seq. The IRR was estimated by comparing the ASV counts per family using a generalized linear model with Poisson regression and log-link function in R at a significance level of p < 0.05. Positive values indicate a significant increase in the detection probability, while negative values indicate a significant decrease in the detection probability.

## Results

### Single-cell diversity & bacterial community composition analysis

The 16S rRNA ASV richness and Shannon diversity of the *T. gravida* single-cell microbiomes was strain specific and impacted by culture condition, but not affected by single-cell DNA processing methods (table S7, Fig. S3). The differences were mainly caused by strain A1 which showed a significantly higher richness and Shannon diversity in the nitrogen- and vitamin-deplete media compared to the full medium (Fig. S4).

Moreover, the 16S rRNA gene ASV community composition of the single-cell *T. gravida*-associated microbiomes differed significantly in response to strain identity and culture conditions (table S8). The genotypic specificity regarding microbiome community composition was reflected on the single-cell level, particularly for highly abundant bacterial groups. This is supported by distinct genotype-specific clustering in the principal component analysis (PCA) (Fig. 2). However, normalized read counts for multiple bacterial genera differed significantly between diatom strains and culture conditions (table S9). Furthermore, no bacterial genus was consistently present across all analyzed single-cell replicates and strains (Fig. 2). Yet, 8 of the 48 detected bacterial genera occurred across all tested strains, but with differences in abundance and frequency between single-cells. Within the fraction of shared bacterial genera, the *Sulfitobacter, Colwellia* and *Roseobacter* clades showed the highest normalized reads and were particularly abundant in strain A1. Moreover, strain A1 shared three bacterial genera exclusively with strain A2 (most importantly *Glaciecola* and *Paraglaciecola*) and four bacterial genera exclusively with strain A5, of which *Polaribacter* was the most abundant genus. In strain A1, eight unique bacterial genera were found, with *Aurantivirga* as the most abundant genus. For strain A2 and A5, ten unique bacterial genera were found each, of which *Lentilitoribacter* showed the highest abundance in strain A2 and *Octadecabacter* clade in strain A5.

**Figure 2:**
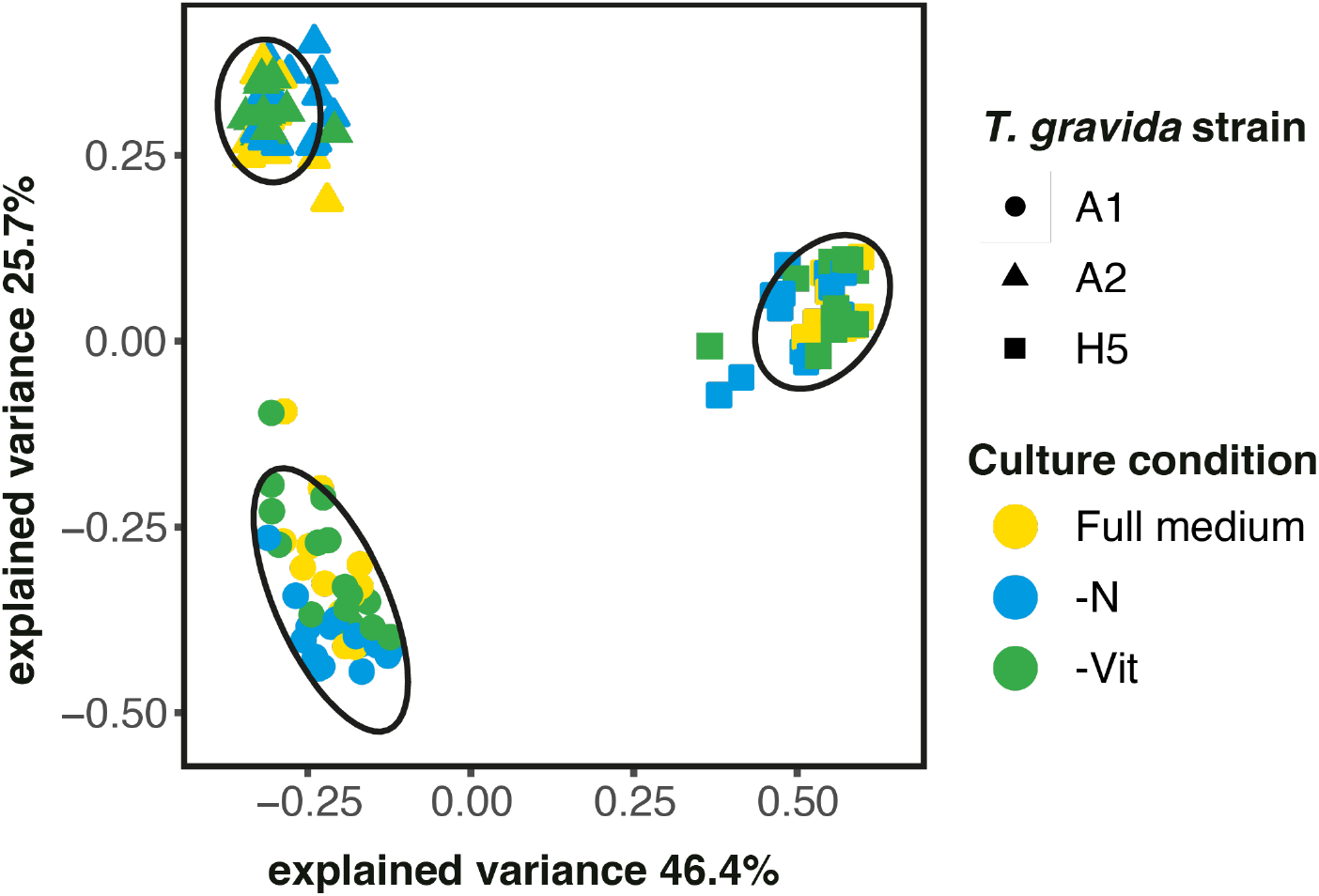
Principal component analysis of Bray-Curtis distances from *T. gravida* single-cell microbiomes in different culture media, including Full K medium (yellow), nitrogen -deplete medium (-N, blue), and vitamin-deplete medium (-Vit, green) based on normalized 16S ASV data. Ellipses (confidence level = 0.95) show clusters of the respective *T. gravida* strain (indicated by shape).

Multiple bacterial groups showed significantly different abundance under specific culture conditions (table S9, Fig. 3). This was evident for *Marinobacter*, which increased in abundance in vitamin-deplete treatments, as well as *Colwellia* showing an increase in abundance under nitrogen-deplete culture conditions.

**Figure 3:**
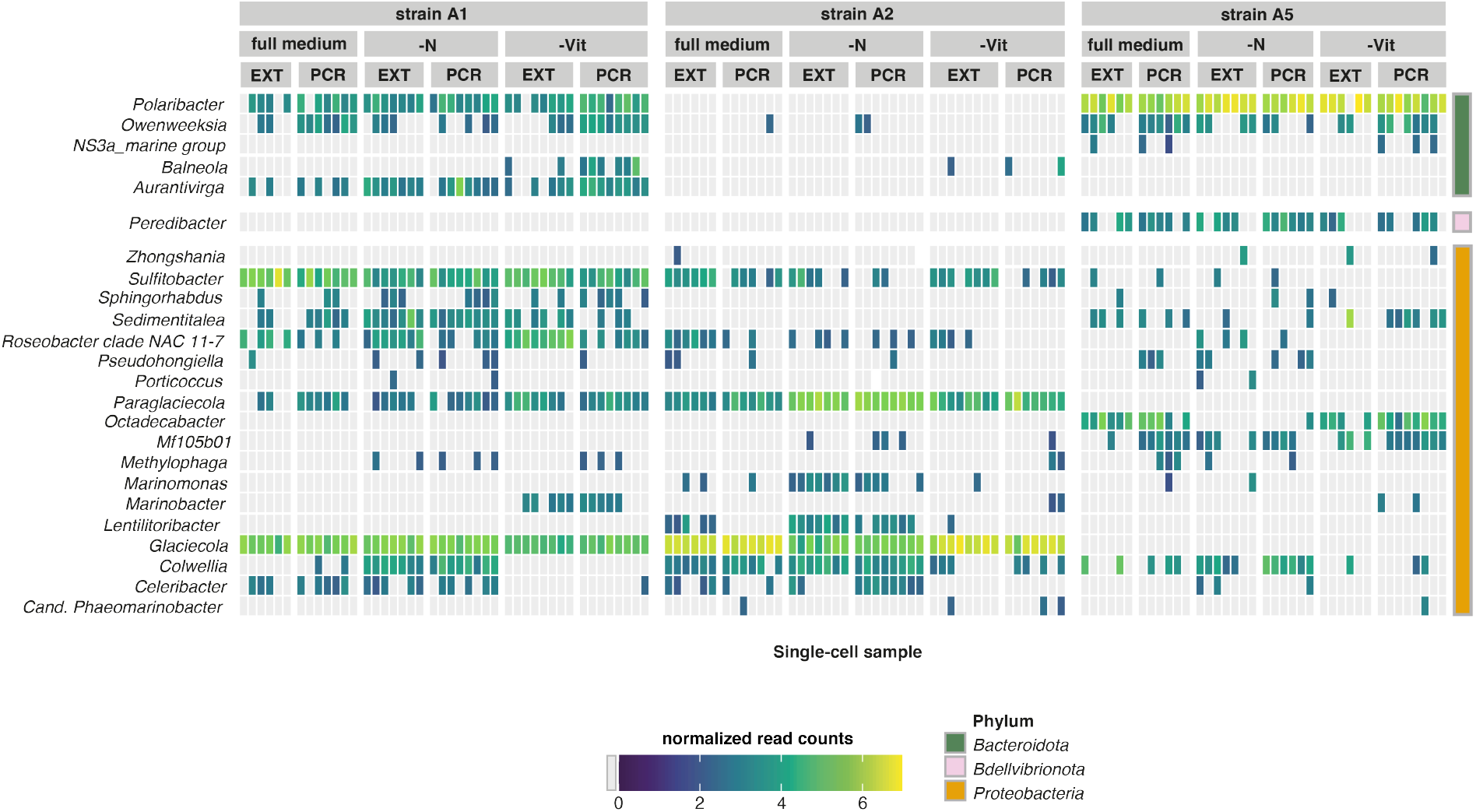
Normalized (subsampled and power-transformed [X^0.25^]) 16S rRNA gene ASV read counts (indicated by continuous color scale, gray = not detected) summarized on the genus level (y-axis) across *T. gravida* single-cell samples (x-axis). *T. gravida* strains and culture conditions are indicated by vertical facets. Genera are ordered by Phylum (displayed as discrete color scale (right). Genera occurring in less than 4 samples were excluded from the plot. Single-cell sample treatments indicated with; full medium = modified K medium, -Vit = Vitamin limited Medium, -N = Nitrogen limited medium, EXT = Single-cell DNA extraction (SC-EXT), PCR = direct Single-Cell PCR (SC-PCR)

### SCHoCO-seq results

The SCHoCO-seq treatment significantly affected the relative proportions of all ASV categories (bacterial, chloroplast, and other ASVs) of the single-cell microbiomes (table 1, Fig. 4). On average, chloroplast ASVs in a given single diatom-cell accounted for 71.4 ± 14.3% of total reads in SC-seq control treatments, but were reduced in the SCHoCO-seq treatment to 17.5 ± 12.8%. Bacterial ASVs, which on average accounted for 25.9 ± 14.7% in the SC-seq control, increased to 71.0 ± 16.7% in the SCHoCO-seq treatment. Random subsampling across ranges of sequencing depth showed that with increasing sequencing depth mostly bacterial 16S ASVs were detected in a theoretical ASV composition in the SCHoCO-seq treatment, whereas increasing sequencing depth in the SC-seq control treatment mainly resulted in more chloroplast ASV reads.

**Table 1:** Results of Welch t-tests on relative proportions of the ASV categories bacteria, chloroplasts and other 16S rRNA gene ASVs as a response to SCHoCO-seq. Bacteria refer to all bacterial ASVs after removal of a) potential contamination and b) ASVs without annotations at least at the order level. Chloroplasts refer to all ASVs annotated as chloroplasts, and ‘Other’ refer to mitochondria-derived ASVs and ASVs with no annotation at least at the order level. Mean proportions of ASV-categories are given in percent ± standard deviation. T- and p-values are reported for the treatment effect for each ASV category. Values marked with an asterisk (*) indicate significant effects (p < 0.05).

|  | % in Cas9 sample | % in No Cas9 sample | p-value | t-value |
| --- | --- | --- | --- | --- |
| Bacteria | 71.0 ± 16.7 | 25.9 ± 14.7 | <0.001 | 15.94 |
| Chloroplast | 17.5 ± 12.8 | 71.4 ± 14.3 | <0.001 | -22.11 |
| Other | 11.5 ± 8.3 | 2.7 ± 2.1 | <0.001 | 8.17 |

**Figure 4:**
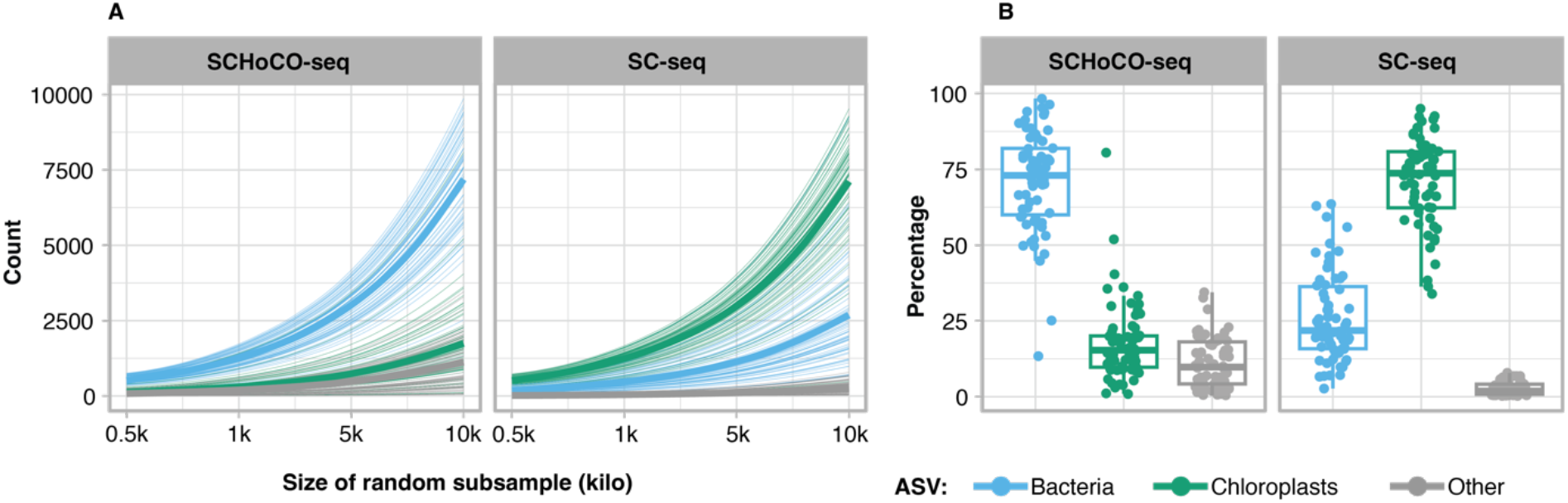
Effect of Cas9 plastidal 16S rRNA gene amplicon digest on total read composition and as a function of a range of assumed sequencing depths. The line plots show the theoretical composition of each sample if it was sequenced at different depths (obtained from random subsamples of the original dataset). Thin lines represent the ASV composition of each of the 62 samples after (SCHoCO-seq) and prior (SC-seq) to the cas9 digest. Bold lines are means obtained from the respective ASV compositions (A). Boxplots represent the percentage of the actual read composition of the ASVs, summed by annotation. ‘Bacteria’ refers to all bacterial ASVs after removal of a) potential contamination and b) ASVs without annotations at least at the order level. ‘Chloroplasts’ refers to all ASVs annotated as chloroplasts, and ‘Others’ refers to mitochondria-derived ASVs and ASVs with no annotation at least at the order level (B).

By increasing the overall proportion of bacterial ASV reads (Fig. 4), SCHoCO-seq significantly increased the probability of detecting several bacterial families compared to SC-seq (Fig. 5A). There was no significant decrease in the detection probability for any bacterial family. The increase in IRR values was particularly pronounced for *Sphingomonadaceae*, where the ASV counts were approximately 16 times higher compared to SC-seq. *Methylophilaceae, Rhizobiaceae, Parvibaculaceae* and *Cryomorphaceae* also showed comparably high IRR values with IRRs > 5. The SCHoCO-seq approach additionally identified the bacterial phylum *Verrucomicrobiota*, which was not covered by the SC-seq control (Fig. 3). Consistently, there was a significant increase in the IRR value for a total of six families within these phyla.

**Figure 5:**
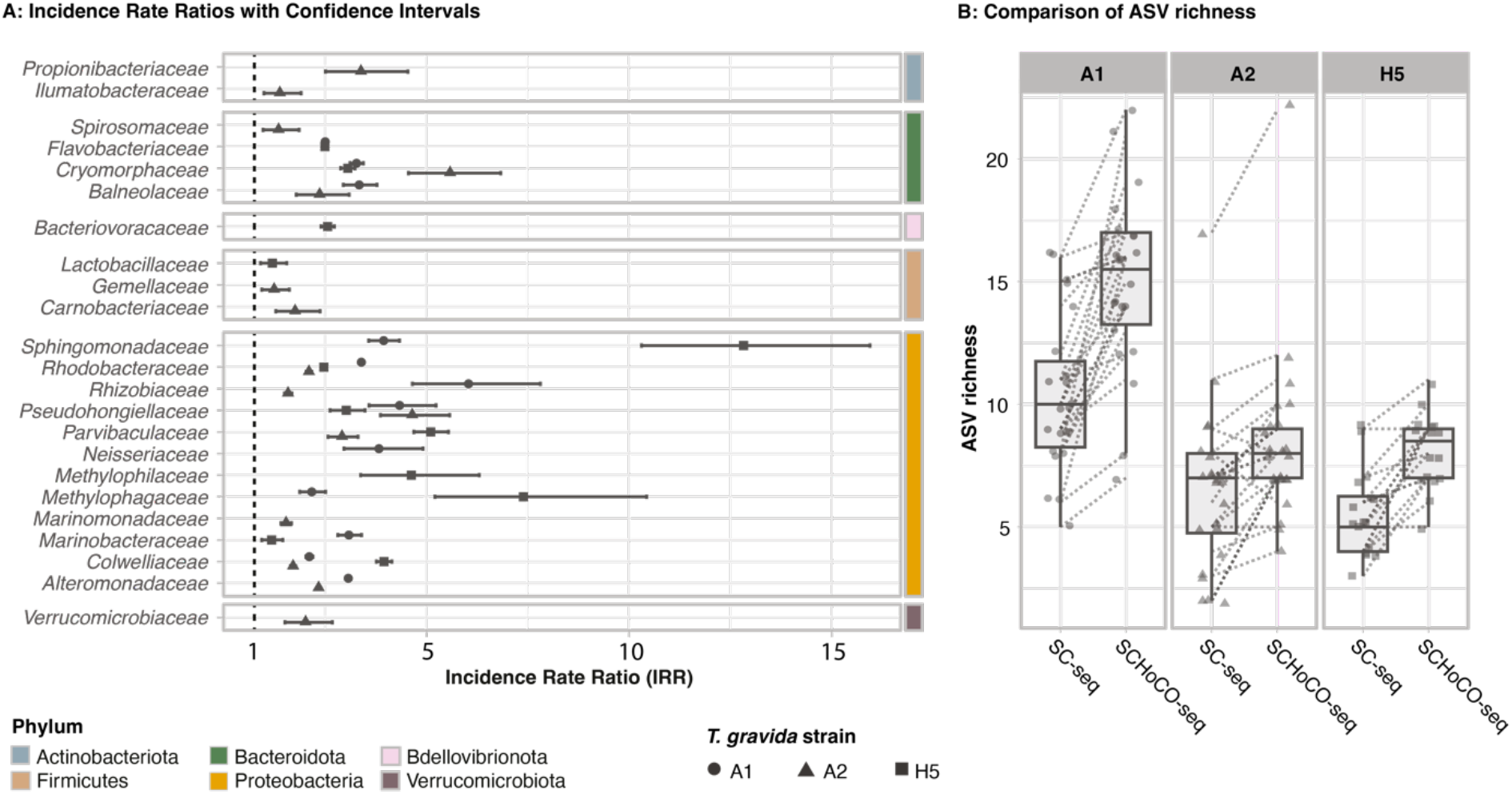
Performance comparison of 62 SCHoCO-seq and SC-seq sample pairs on A: the differences in the probability of detecting a given bacterial family level based on ASV counts per sample and B) the actual total ASV richness. In A, bacterial families for which a significant change in the Incidence Rate Ratio could be detected are shown, with error bars representing the 95% confidence interval. The dotted line marks the boundary between a decrease in IRR (<1) and an increase in IRR (>1).

In line, the ANOVA results indicated a significant effect of SCHoCO-seq treatment on bacterial ASV richness (Fig. 5B, table S10). Single-cell samples without Cas9 digest (SC-seq) had a mean ASV richness of 10.4 ± 3.2 (strain A1), 6.5 ± 3.3 (strain A2), and 5.6 ± 1.8 (strain A5). In SCHoCO-seq treatments, richness increased to 15.0 ± 3.7 (strain A1), 8.4 ± 3.4 (strain A2), and 8.1 ± 1.5 (strain A5), corresponding to a richness increase of 51%, 53%, and 56%, respectively.

### In-silico gRNA specificity

The selected gRNA and the search for corresponding crRNA target sequences in the full PhytoREF databases resulted in a perfect match (100%) with 969 of the 1068 available diatom sequences (∼90%), spanning various taxonomic lineages (Fig. 6A). Of 99 sequences not matching the crRNA, 85 had at least no genus-level annotation, rendering their taxonomic identification elusive. Identification of potential Cas9 off-targets based on the 16S rRNA gene SSU sequences obtained from the SILVA V138 database resulted in 100% matches to only 644 sequences annotated as chloroplast and one mitochondrial hit. However, there were no other matches to any of the 451,459 16S rRNA gene SSU sequences, indicating no potential Cas9 off-targets for the selected gRNA. Low-probability potential off-targets (i.e., considering a gRNA sequence match of 96%, (Hsu et al., 2013)) were identified for 117 16S rRNA gene SSUs, representing only a fraction of 0.025% of the sequence representatives in the SILVA V138 database. The evaluation of the phylogenetic range of the diatom chloroplast sequences based on the crRNA target sequence and the outer and inner primers used for SCHoCO-seq resulted in 62 matches, spanning at least 3 families and 12 orders (Fig. 6B). It should be noted that this relatively low number compared to the gRNA matches is due to the fact that only ∼20% of sequences in PhytoREF are long enough to fully cover both primer binding sites.

**Figure 6:**
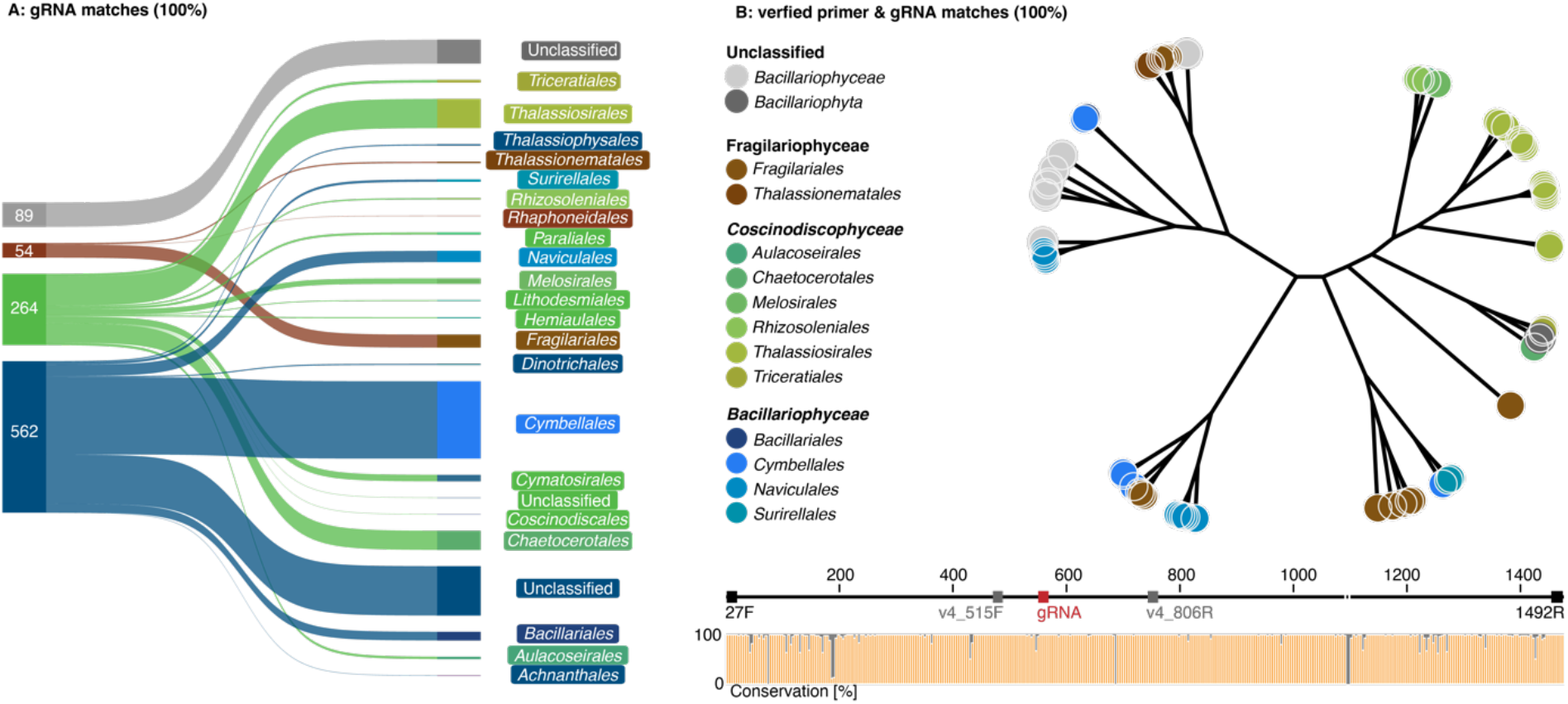
Coverage of taxonomic lineages with perfect gRNA sequence matches (A) and perfect matches with all primer pairs used in this study (B). Numbers in (A) refer to the gRNA chloroplast sequence matches for each diatom family. The bottom panel in (B) shows the primer binding sites and nucleotide conservation of 62 diatom chloroplast sequences; the corresponding unrooted phylogenetic tree shows the order-level annotations as indicated in the PhytoREF database.

## Discussion

This study demonstrated that diatom microbiomes under investigation were mostly strain-specific and that the different micro-/macronutrient limitations treatments modulated the abundance of single microbiome members. The SCHoCO-seq treatment enabled in depth assessment of the microbial diversity in algal microbiomes on single-cell host level by reducing chloroplast ASV reads while increasing sequencing depth for bacterial ASVs. Consequently, SCHoCO-seq led to increased microbiome richness and detected rare microbial taxa which were not found with standard 16S rRNA gene single-cell metabarcoding. Our *in-silico* analyses identified a 100% match to 969 of the potential 1068 crRNA target sequences in PhytoREF databases, suggesting the newly developed protocol to be applicable to a wide range of diatom species with no apparent 16S rRNA gene off-target effects.

With respect to genotype-specific microbiomes of *T. gravida*, strain A1 and A2 were more similar to each other than to strain A5. Despite differing dates of isolation, variations in microbiome composition were most likely explained by the biogeographic distance of their sampling origin (table S1), as microbiomes of laboratory cultures have been shown to be stable over long periods of time (Barreto Filho et al., 2021). Yet, although strain A1 and A2 originated from the same location, their microbiomes still showed differences in richness and diversity, highlighting the necessity for a single-cell perspective when profiling natural phytoplankton microbiomes.

Previous studies discussed the possibility of a core microbiome, i.e., a fraction of microbiome bacteria consistently present across diatom genotypes (Behringer et al., 2018, Monnich et al., 2020). However, the probability to detect a core microbiome likely decreases with increasing sampling size, biogeographic distance and region. For example, Ahern et al. (2021) found no shared ASVs across 85 *Thalassiosira rotula* strains from various ocean provinces. Yet, for the *T. gravida* strains in this study, a shared core community across the studied genotypes was indeed found, although not consistently across all single-cell replicates. This finding reflects the diatom’s Arctic origin, where host-microbe associations are more stable due to co-evolution under hard selection compared to the soft selection in temperate regions (Malmstrom et al., 2007). Remarkably, the shared core microbial community consisted of the genera *Colwellia, Roseobacter clade NAC11-7 lineage* as well as *Sulfitobacter*, also belonging to the *Roseobacter* group. Particularly the *Sulfitobacter* genus evolved complex mutualistic-symbiotic interactions with phytoplankton of which the underlying mechanisms are comparably well understood and include bacteria-derived provision of auxin and ammonium for their host (Amin et al., 2015, Segev et al., 2016), as well as specific antibiotics to protect the holobiont from parasitic bacteria (Beiralas et al., 2023). In exchange, *Sulfitobacter* receive various forms of exudated organic carbon from the diatom host, as well as dimethylsulfoniopropionate (DMSP) for sulfur oxidation, and amino acids like tryptophan and taurine as organic nitrogen source (Amin et al., 2015, Segev et al., 2016, Yang et al., 2021).

Moreover, the *T. gravida* microbiomes responded to nitrogen- or vitamin-limited culture conditions with compositional changes of few specific bacterial genera. Considering that the nitrogen- and vitamin-limited cultures were inoculated from a full medium culture, the fact that certain bacterial groups were only retrieved in the limitation treatments suggests that these rare microbiome taxa may become abundant only during abiotic stress. This is exemplified for strain A1, where *Marinobacter* increased in abundance in the vitamin-limited treatments. *Marinobacter* are known for their ability to produce siderophores, potentially providing iron to their host (Amin et al., 2009, Butler et al., 2021). *Marinobacter* have also been associated with potential vitamin supply for the marine dinoflagellate *Lingulodinium polyedrum* (Cruz-Lopez and Maske, 2016). For the genus *Balneola*, an increase in the nutrient-limited treatments was observed for *T. gravida* strain A1. However, these increases were not necessarily due to a mutualistic interaction but may also indicate opportunistic mechanisms. In fact, Zhu et al. (Zhu et al., 2021) hypothesized that *Balneola* has the ability to degrade specific organic compounds released by plants and potentially also phytoplankton upon encounter of nutrient stress. Moreover, diatoms have been shown to regulate their microbiome by target-specific secondary metabolites like rosmarinic acid or azelaic acid (Shibl et al., 2020), yet whether this was the case in our experiments remains elusive.

Studying microbiome diversity in laboratory cultures is influenced not only by the fact that most heterotrophic bacteria cannot be cultured (Vartoukian et al., 2010), but also by the dynamics of the microbiome community itself. Certain culture conditions may favor the growth of specific bacterial groups, thus underestimating bacterial richness. Conversely, shifts in evenness may allow the detection of bacterial groups that perform essential services for their host and represent an insurance for the holobiont in case of perturbations, while others remain undetected. This highlights the importance of considering microbiome community dynamics and the potential for different culture conditions to reveal distinct aspects of the microbiome’s diversity and functional roles. Additionally, considering the isolation bias in terms of species- and even genotype-specificity of microbiome composition (Baker and Kemp, 2014, Ahern et al., 2021), our findings underline the necessity for (semi-) quantitative *in-situ* single-cell data on microbiome composition.

The bacterial richness in *T. gravida* microbiomes obtained in this study at the single-cell level by SCHoCO-seq agrees with richness data obtained from bulk diatom samples ranging from 1 to 125 ASVs or OTUs, respectively (Baker and Kemp, 2014, Behringer et al., 2018, Ahern et al., 2021, Sison-Mangus et al., 2014). To recover the full bacterial community richness on a given single diatom cell (preceding the SCHoCO-seq protocol), both the SC-PCR and SC-EXT methods developed in this study have been equally accurate as reflected by insignificant statistical test results. Both single-cell methods are therefore applicable depending on the research objective. For example, the SC-EXT approach offers the advantage of enabling multiple PCR reactions for further genomic DNA based investigations, such as metagenomics or host identification via 18S sequencing, and the long-term storage of the sample (Hallmaier-Wacker et al., 2018).

The SCHoCO-seq approach is a promising strategy to accurately analyze diatom host-associated microbiome ASVs. This is largely due to the reduction of chloroplast reads, which in turn boost the detection and resolution of the microbiome richness. Furthermore, our subsampling approach showed that the increase of microbiome ASVs over various sequencing depths by SCHoCO-seq was higher than the corresponding increase in chloroplast ASVs (Fig. 4). Importantly, this pattern was the opposite without Cas9 treatment, suggesting that merely increasing the sequencing depth is not an efficient strategy to recover rare microbiome ASVs. Therein, SCHoCO-seq is particularly valuable in resolving the bacterial diversity of diatom microbiomes that harbor low bacterial abundance in their phycosphere. Thus, especially due to its potential to reveal higher community richness and identify potentially rare bacterial taxa in the phycosphere, SCHoCO-seq has the potential to become the preferred method in diatom microbiome diversity research. Indeed, by identifying rare bacterial taxa, taxon shifts, gains and losses, the overall genetic diversity in diatom microbiomes can be more accurately assessed, thereby increasing our knowledge of microbiome community assembly (Monnich et al., 2020), regulation and modulation. As microbiome genetic diversity and variation are aligned to selection (Baltar et al., 2019), these capabilities are particularly relevant under current global change conditions. SCHoCO-seq therefore bears potential to assess and distinguish diatom microbiome responses to environmental change, e.g., plasticity, resilience, functional redundancy and dysbiosis of diatom holobionts (Baltar et al., 2019, Graham et al., 2016).

## Conclusion

This study revealed a) genotype-specific diatom microbiomes identified through single-cell isolation and b) that diatom holobionts respond to macro- and micro-nutrient limitations with compositional changes of their microbiome. To increase the discovery of rare bacterial taxa as part of diatom microbiomes, we have developed SCHoCO-seq which increases the bacterial biodiversity (richness) by reducing the chloroplast 16S ASVs. Thus, this novel single-cell microbiome analysis may serve as an important steppingstone towards *in-situ* single-cell microbiome analysis. The latter will help to understand how phytoplankton-associated microbiomes function in natural systems. In addition, the method presented here for reducing host DNA contamination with the aid of Cas9 can be applied to various other host-microbiome systems by adapting the sequence of the guide RNA.

## Acknowledgments

The authors thank the working group for their support.

## Author Contributions

Ruben Schulte-Hillen and Jakob Giesler contributed equally to this work and share first authorship. For the purposes of applications, proposals and outreach, either author may list themselves first.

## Data availability

All raw sequence data used in this study will be made available upon publication.

