## Supplementary Material for "A CRISPR-Cas9 assisted analysis of single-cell microbiomes for identifying rare bacterial taxa in phycospheres of diatoms"

Ruben Schulte-Hillen*[1,2,3], Jakob K. Giesler*[1,4], Thomas Mock[5], Nigel Belshaw[5], Uwe John[1], Tilmann Harder[1,4], Nancy Kühne[1], Stefan Neuhaus[1], and Sylke Wohlrab[1,6]

* These authors contributed equally to this work and share first authorship.

[1)] *Alfred Wegener Institute, Helmholtz Centre for Polar and Marine Research, Bremerhaven Germany*

[2)] *Albert-Ludwigs-Universität Freiburg, Freiburg, Germany*

[3)] *Max Planck Institute for Marine Microbiology, Bremen, Germany*

[4)] *Universität Bremen, Bremen, Germany*

[5)] *University of East Anglia, School of Environmental Sciences, Norwich, UK*

[6)] *Helmholtz Institute for Functional Marine Biology at the University of Oldenburg (HIFMB), Oldenburg, Germany*

**Table S5.1:** Detailed information about *Thalassiosira gravida* strains used in this study.

| name | NORCCA strain number | isolation date | lat | lon |
|---|---|---|---|---|
| A1 | UIO478 | 20.08.17 | 83.33 | 29.29 |
| A2 | UIO483 | 05.09.17 | 83.15 | 31.46 |
| A5 | UIO448 | 12.09.17 | 79.99 | 14.97 |

**Table S5.2:** Recipe for full (modified) K – seawater medium. Final concentrations refer to the addition of the respective components and do not consider potential background concentration in seawater.

| Full (modified) K - medium | |
|---|---|
| Component | Final concentration |
| Enrichments | |
| $NaNO_3$ | $8.82 \times 10^{-4}$ M |
| $NaH_2PO_4 \times H_2O$ | $1.00 \times 10^{-5}$ M |
| $Na_2SiO_3 \times 9\ H_2O$ | $1,06 \times 10^{-4}$ M |
| $H_2SeO_3$ | $1.00 \times 10^{-8}$ M |
| Tris Base | $1.00 \times 10^{-3}$ M |
| Supplements (trace metals) | |
| $Na_2EDTA \times 2\ H_2O$ | $1.12 \times 10^{-4}$ M |
| $FeCl_3 \times 6\ H_2O$ | $1.17 \times 10^{-5}$ M |
| $Na_2MoO_4 \times 2H_2O$ | $2.60 \times 10^{-8}$ M |
| $ZnSO_4 \times 7\ H_2O$ | $7.65 \times 10^{-8}$ M |
| $CoCl_2 \times 6\ H_2O$ | $4.20 \times 10^{-8}$ M |
| $MnCl_2 \times 4\ H_2O$ | $9.10 \times 10^{-7}$ M |
| $CuSO_4 \times 5\ H_2O$ | $1.96 \times 10^{-8}$ M |
| Supplements (vitamins) | |
| Vitamin B12 | $3.69 \times 10^{-10}$ M |
| Biotin | $2.05 \times 10^{-9}$ M |
| Thiamin HCl | $2.96 \times 10^{-7}$ M |

**Table S5.3:** Recipe for nitrogen limited (modified) K – seawater medium. Final concentrations refer to the addition of the respective components and do not include background concentration in the seawater. By omitting the addition of inorganic nitrogen sources to the medium, the final nitrogen concentration is limited to the background concentration in the seawater (~ 5 µM $NaNO_3$; ~ 5 µM $NH_4$).

| Nitrogen limited (modified) K - Medium | |
|---|---|
| Component | Final concentration |
| **Enrichments** | |
| $NaNO_3$ | - |
| $NaH_2PO_4$ x $H_2O$ | 1.00 x $10^{-5}$ M |
| $Na_2SiO_3$ x 9 $H_2O$ | 1,06 x $10^{-4}$ M |
| $H_2SeO_3$ | 1.00 x $10^{-8}$ M |
| Tris Base | 1.00 x$10^{-3}$ M |
| **Supplements (trace metals)** | |
| $Na_2EDTA$ x 2 $H_2O$ | 1.12 x $10^{-4}$ M |
| $FeCl_3$ x 6 $H_2O$ | 1.17 x $10^{-5}$ M |
| $Na_2MoO_4$ x $2H_2O$ | 2.60 x $10^{-8}$ M |
| $ZnSO_4$ x 7 $H_2O$ | 7.65 x $10^{-8}$ M |
| $CoCl_2$ x 6 $H_2O$ | 4.20 x $10^{-8}$ M |
| $MnCl_2$ x 4 $H_2O$ | 9.10 x $10^{-7}$ M |
| $CuSO_4$ x 5 $H_2O$ | 1.96 x $10^{-8}$ M |
| **Supplements (vitamins)** | |
| Vitamin B12 | 3.69 x $10^{-10}$ M |
| Biotin | 2.05 x $10^{-9}$ M |
| Thiamin HCl | 2.96 x $10^{-7}$ M |

**Table S5.4:** Recipe for vitamin limited (modified) K – seawater medium. Final concentrations refer to the addition of the respective components and do not include background concentration in the seawater. However, due to autoclaving, vitamin concentrations in the seawater were negligible.

| Vitamin limited (modified) K - Medium | |
|---|---|
| Component | Final concentration |
| Enrichments | |
| $NaNO_3$ | $8.82 \times 10^{-4}$ M |
| $NaH_2PO_4 \times H_2O$ | $1.00 \times 10^{-5}$ M |
| $Na_2SiO_3 \times 9\ H_2O$ | $1{,}06 \times 10^{-4}$ M |
| $H_2SeO_3$ | $1.00 \times 10^{-8}$ M |
| Tris Base | $1.00 \times 10^{-3}$ M |
| Supplements (trace metals) | |
| $Na_2EDTA \times 2\ H_2O$ | $1.12 \times 10^{-4}$ M |
| $FeCl_3 \times 6\ H_2O$ | $1.17 \times 10^{-5}$ M |
| $Na_2MoO_4 \times 2H_2O$ | $2.60 \times 10^{-8}$ M |
| $ZnSO_4 \times 7\ H_2O$ | $7.65 \times 10^{-8}$ M |
| $CoCl_2 \times 6\ H_2O$ | $4.20 \times 10^{-8}$ M |
| $MnCl_2 \times 4\ H_2O$ | $9.10 \times 10^{-7}$ M |
| $CuSO_4 \times 5\ H_2O$ | $1.96 \times 10^{-8}$ M |
| Supplements (vitamins) | |
| Vitamin B12 | - |
| Biotin | - |
| Thiamin HCl | - |

**Table S5.5:** Modified DNA Extraction Protocol based on DNA MasterPure Complete DNA & RNA Purification Kit for fluid samples.

| Step | Procedure |
|---|---|
| 1 | Thaw frozen samples (sample in 30 µL of 2xT&C Lysis buffer) at 50°C |
| 2 | Add 30 µL ultrapure $H_2O$ |
| 3 | Add 30 µL mastermix (prepared as 30 µL 2xT&C Lysis Buffer + 0.2 µL Protease K per sample) at room temperature |
| 4 | Incubate for 15 min at 65°C and 1,000 rpm; cool for 5 min on ice |
| 5 | Add 45 µL MPC protein precipitation reagent; vortex for 10 s |
| | Centrifuge at 10,000 x g, 4°C for 10 min; transfer supernatant to new 1.5 mL reaction vile |
| 6 | Add 150 µL isopropanol and 0.7 µL Pellet Paint NF Co-Precipitant to each sample |
| 7 | Centrifuge at 15,000 x g, 4°C for 10 min; discard supernatant |
| 8 | Wash DNA pellet twice with 150 µL of 70% ethanol |
| 9 | Dissolve DNA pellet in 12.5 µL TE buffer |
| 10 | Store at -20°C |

**Table S5.6:** List of common contaminants according to Sheik et al. (2018) and other non-marine bacteria that were excluded from analysis (if present).

| **Contaminant (Genus)** | | | |
|---|---|---|---|
| *Abiotrophia* | *Cloacibacterium* | *Klebsiella* | *Polaromonas* |
| *Acidobacteria_Gp2* | *Comamonas* | *Kocuria* | *Prevotella* |
| *Acidovorax* | *Corynebacterium* | *Lactobacillus* | *Prevotella_7* |
| *Acinetobacter* | *Craurococcus* | *Lawsonella* | *Propionibacterium* |
| *Acinetobactera* | *Cupriavidus* | *Leptothrix* | *Pseudomonas* |
| *Actinomyces* | *Curtobacterium* | *Limnobacter* | *Pseudoxanthomonas* |
| *Aerococcus* | *Curvibacter* | *Massilia* | *Psychrobacter* |
| *Aeromicrobium* | *Cutibacterium* | *Mesorhizobium* | *Ralstonia* |
| *Afipia* | *Deinococcus* | *Methylobacterium* | *Rhizobium* |
| *Alloprevotella* | *Delftiae* | *Methylophilus* | *Rhodococcus* |
| *Anaerococcus* | *Devosia* | *Methyloversatilis* | *Roseomonas* |
| *Aquabacterium* | *Dietzia* | *Microbacterium* | *Rothia* |
| *Arthrobacter* | *Duganella* | *Micrococcus* | *Ruminococcus* |
| *Asticcacaulis* | *Dyadobacter* | *Microlunatus* | *Schlegelella* |
| *Aurantimonas* | *Enhydrobacter* | *Neisseria* | *Sphingobium* |
| *Azoarcus* | *Enterobacter* | *Nevskia* | *Sphingomonas* |
| *Azospira* | *Escherichia* | *Niastella* | *Sphingopyxis* |
| *Bacillus* | *Escherichia-Shigella* | *Novosphingobium* | *Staphylococcus* |
| *Beijerinckia* | *Escherichia/Shigella* | *Ochrobactrum* | *Stenotrophomonas* |
| *Beutenbergia* | *Facklamia* | *Olivibacter* | *Streptococcus* |
| *Bosea* | *Finegoldia* | *Oxalobacter* | *Sulfuritalea* |
| *Bradyrhizobium* | *Flavobacterium* | *Paenibacillus* | *Tsukamurella* |
| *Brevibacillus* | *Fusobacterium* | *Parabacteroides* | *Turicella* |
| *Brevibacterium* | *Geodermatophilus* | *Paracoccus* | *Undibacterium* |
| *Brevundimonas* | *Haemophilus* | *Pasteurella* | *Variovorax* |
| *Brochothrix* | *Herbaspirillum* | *Patulibacter* | *Veillonella* |
| *Burkholderia* | *Hoefleae* | *Pedobacter* | *Wautersiella* |
| *Capnocytophaga* | *Hydrotalea* | *Pedomicrobium* | *Xanthomonas* |
| *Cardiobacterium* | *Janibacter* | *Pelomonas* | |
| *Caulobacter* | *Janthinobacterium* | *Peptoniphilus* | |
| *Chryseobacterium* | *Kingella* | *Phyllobacterium* | |

**Table S5.7:** ANOVA results testing differences in bacterial richness and Shannon diversity of *T. gravida* single cell microbiomes as a response of strain identity, culture condition and DNA processing method. F and p-values are reported for each effect. Values marked with an asterisk (*) indicate significant effects ($p < 0.05$).

| | | Richness | | | Shannon | | |
|---|---|---|---|---|---|---|---|
| *Effect* | *Df* | *F* | *p* | | *F* | *p* | |
| Strain | 2 | 27.525 | <0.001 | * | 27.905 | <0.001 | * |
| Condition | 2 | 4.965 | 0.008 | * | 6.283 | 0.003 | * |
| Method | 1 | 0.476 | 0.492 | | 0.488 | 0.486 | |
| Strain*Condition | 4 | 3.424 | 0.011 | * | 4.108 | 0.004 | * |

**Table S5.8:** PERMANOVA results testing differences in bacterial community composition of *T. gravida* single cell microbiomes as a response of strain identity and culture condition. F and p-values are reported for each effect. Values marked with an asterisk (*) indicate significant effects ($p < 0.05$).

| | | microbiome composition | | | |
|---|---|---|---|---|---|
| *Effect* | *Df* | $R^2$ | *F* | *p* | |
| Strain | 2 | 0.700 | 199.9 | <0.001 | * |
| Condition | 2 | 0.042 | 12.1 | <0.001 | * |
| Strain*Condition | 4 | 0.048 | 6.9 | <0.001 | * |

**Table S5.9:** Multiple ANOVA results on normalized ASV read counts summarized on the genus level as a response to strain identity, culture condition and their respective interactive effect. Combinations of bacterial genera and respective effects marked with an asterisk indicate significant effects (significance code: * = p < 0.05 & > 0.01; ** = p < 0.01 & > 0.001; *** = p < 0.001).

| Genus | strain | condition | strain*condition |
|---|---|---|---|
| *Adhaeribacter* | | | |
| *Alloiococcus* | | | |
| *Amaricoccus* | | | |
| *Aurantivirga* | *** | ** | *** |
| *Balneola* | ** | *** | *** |
| *Candidatus_Phaeomarinobacter* | | | |
| *Celeribacter* | *** | *** | * |
| *Colwellia* | *** | *** | ** |
| *Croceibacter* | | | |
| *Fimbriiglobus* | | | |
| *Friedmanniella* | | | |
| *Gemella* | | | |
| *Glaciecola* | *** | *** | *** |
| *Granulicatella* | | | |
| *Ilumatobacter* | | | |
| *Lentilitoribacter* | *** | *** | *** |
| *Luteimonas* | | | |
| *Maribacter* | | | |
| *Marinobacter* | *** | *** | *** |
| *Marinomonas* | *** | *** | *** |
| *Methylobacterium-Methylorubrum* | | | |
| *Methylophaga* | | | * |
| *Mf105b01* | *** | | |
| *NS3a_marine_group* | *** | | |
| *Octadecabacter* | *** | *** | *** |
| *Oleiphilus* | * | | |
| *Owenweeksia* | *** | | |
| *Pacificibacter* | | | |
| *Paraglaciecola* | *** | *** | *** |
| *Peredibacter* | *** | | |
| *Phaeobacter* | | | |
| *Polaribacter* | *** | | |
| *Porticoccus* | | * | |
| *Pseudohongiella* | | ** | |
| *Reichenbachiella* | | | |
| *Romboutsia* | | | |
| *Roseivirga* | | | * |
| *Roseobacter clade NAC11-7 lineage* | *** | | ** |
| *Sedimentitalea* | *** | | *** |
| *Segetibacter* | | | |
| *Sphingorhabdus* | *** | | |
| *Spirosoma* | | | |
| *Sulfitobacter* | *** | ** | |
| *Thalassospira* | | | * |
| *Treponema* | | | |
| *Verticiella* | | | |
| *Zhongshania* | | | |

**Table S5.10:** Results of ANOVA testing the effects of SCHoCO-seq treatment on bacterial ASV richness for microbiomes of *T. gravida* single cells. Single cell ID was treated as random factor. Degrees of freedom (df), mean squares (Mean Sq), F-, and p-values are given. Values marked with an asterisk (*) indicate significant effects (p < 0.05).

| | | | Richness | | |
|---|---|---|---|---|---|
| Main Effect | Df | Mean Sq | *F* | *p* | |
| SCHoCO-seq | 1 | 282.01 | 81.34 | < 0.001 | * |
| Random Effect | | | | | |
| Single-Cell ID | 61 | 29.68 | - | - | |

**Fig. S5.1:** Gel-electrophoresis of 16S rDNA Amplicons. Single Cell Samples from Strain A1 (top), A2 (middle) and A5 (bottom). Eight single cells from full medium (left), from vitamin depleted medium (middle) and from nitrogen depleted medium (right). Culture blanks (culture samples without diatom cells) on the left of each row for all three strains. Size of Amplicons is around 1.5 kb according to the 1 kb DNA Ladder (Gold Biotechnology Inc, USA).

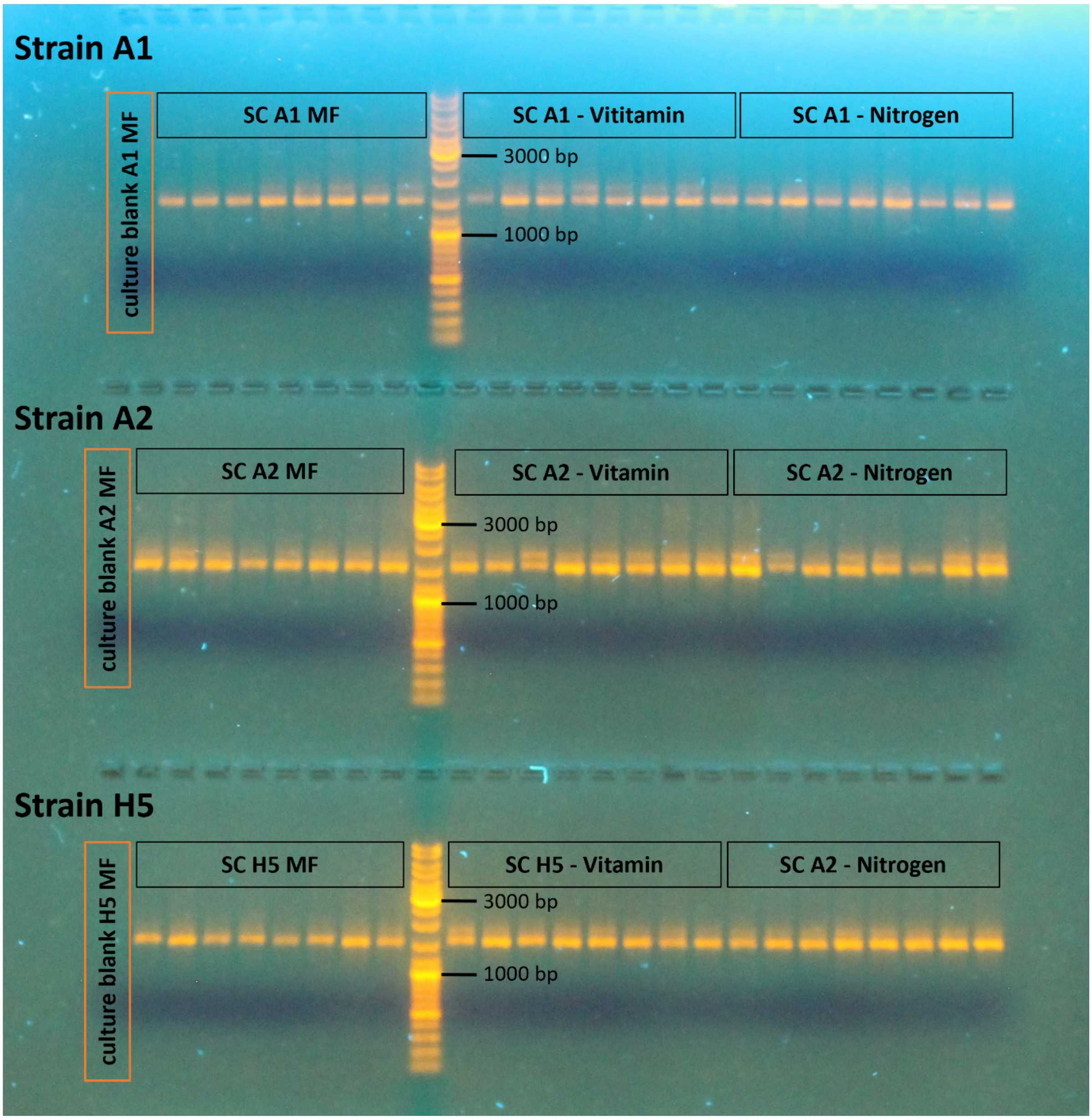

**Fig. S5.2:** Design of gRNA from four oligonucleotides. Primer F and Primer R are needed to generate full length gDNA copies during PCR. The Specific oligo #7 contains the target binding sequence (red) and is linked to the universal oligo containing the Cas9 Nuclease binding site by the linker region (green).

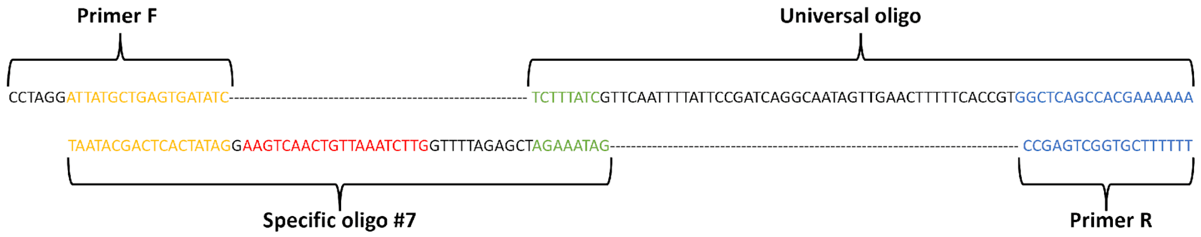


**Fig. S5.3:** Bacterial richness (right) and Shannon diversity (left) of *T. gravida* single cell microbiomes across two different DNA processing methods (i.e., extracted DNA vs. direct PCR).

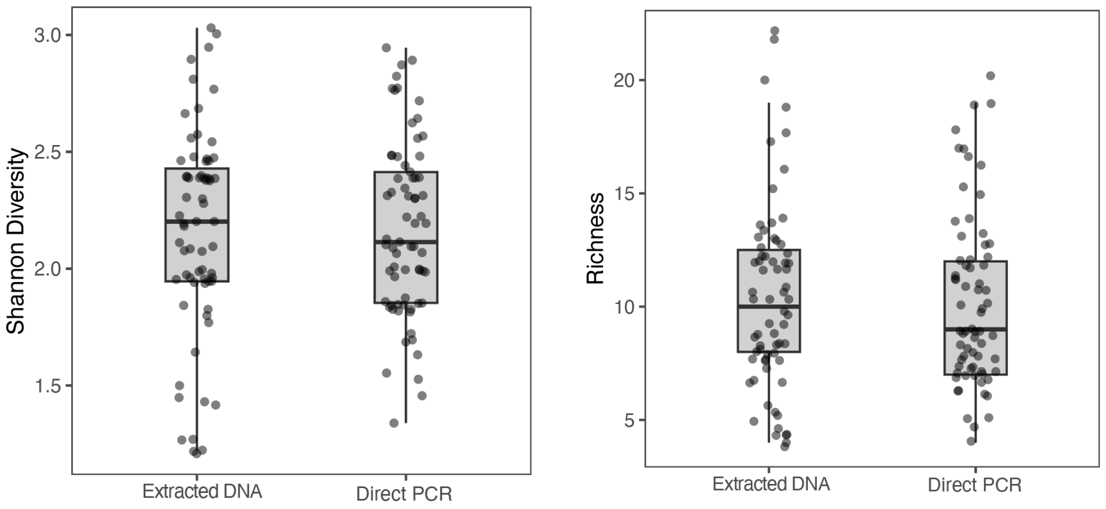

**Fig. S5.4:** Bacterial richness (top row) and Shannon diversity (bottom row) of T. *gravida* single cell microbiomes across strains and culture conditions represented as horizontal facets.

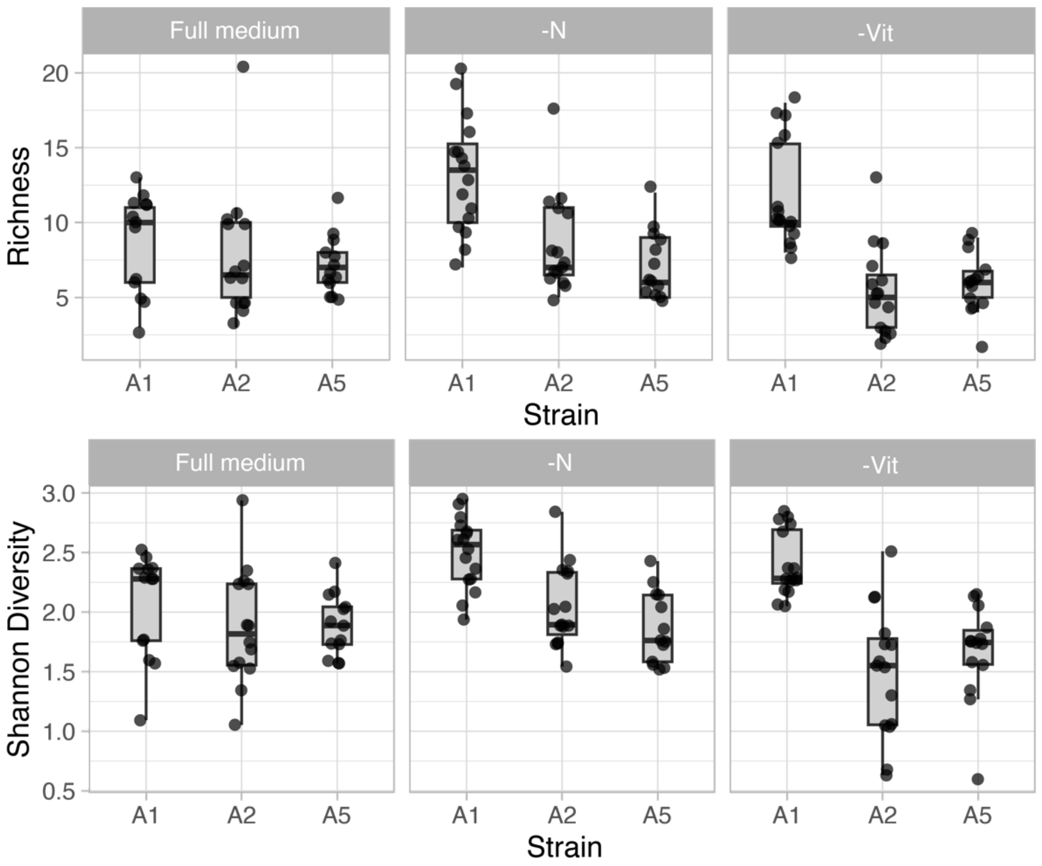
